# Assessing chromatin accessibility in maize using ATAC-seq

**DOI:** 10.1101/526079

**Authors:** Yi-Jing Lee, Pearl Chang, Jui-Hsien Lu, Pao-Yang Chen, Chung-Ju Rachel Wang

## Abstract

**Background:** Maize is an important crop that has a complex genome. A better understanding of maize chromatin architecture provides great opportunities for crop improvement, because chromatin accessibility influences gene expression, thereby affecting agricultural traits. The newly developed method for chromatin profiling, Assay for Transposase Accessible Chromatin with high-throughput sequencing (ATAC-seq), has been developed to investigate chromatin accessibility.

**Result:** We adapt this method by testing parameters of several key steps and generate the first ATAC-seq protocol for maize. We demonstrate that purification of maize nuclei to eliminate organelles can be achieved without the need for cell sorting, and that only a standard bench-top centrifuge is required for sample preparation. Finally, our sequence analyses confirm that our protocol of ATAC-seq can be successfully used to assess the chromatin landscape in maize.

**Conclusion:** The ATAC-seq provides a useful technique to study the chromatin accessibility. Given the parameters tested in our study, it can be a simple and practical method for maize and may be a foundation for similar studies in other crop species.

## Introduction

In eukaryotes, genomic DNA is packaged with histone proteins, forming nucleosomes that constitute the structural basis of chromatin (1). The density of nucleosomes determines chromatin compaction and DNA accessibility, which regulate various cellular and chromosomal functions (2). Genomic regions with dense nucleosomes are tightly packed (i.e., “closed”), whereas nucleosome-depleted regions with exposed DNA are more accessible (i.e., “open”), so the dynamic structures of chromatin provide different levels of availability of DNA binding sites in regulatory regions to transcription factors (3). It is now evident that chromatin structure is intimately linked to the activity of underlying genes, thus influencing proper development and the ability to adapt to an ever-changing environment.

Over the last decade, the development of a wide range of methods that utilize nuclease enzymes such MNase and DNase I to target open DNA regions, combined with next generation sequencing (NGS), has enabled genome-wide investigations of chromatin accessibility (4, 5). For example, MNase digests the linker DNA between nucleosomes, so subsequent NGS reads largely represent the footprints of nucleosomes, which can be used to assess nucleosome occupancy (6). Another method utilizes an optimized concentration of non-specific endonuclease DNase I that generates DNA fragments by liberating open chromatin regions. The resulting NGS reads are characterized as DNase I hypersensitive sites (DHSs), which represent accessible regions of chromatin (7). These methods have been implemented in a number of organisms, ranging from yeast to plants and human. However, they usually require millions of cells, empirical enzymatic titrations, as well as several purification steps, rendering them challenging for reproducible evaluation of chromatin status and consequently they are infeasible for some cell types (8).

An improved method for identifying accessible chromatin is Assay for Transposase-Accessible Chromatin with high-throughput sequencing (ATAC-seq) (9, 10). This method takes advantage of the engineered prokaryotic transposase Tn5 that cleaves DNA in open chromatin regions and simultaneously integrates adapters into cleavage sites *in vivo*. Due to the integrated adapters loaded by Tn5 transposase, the ligation and additional purification steps prior to sequencing necessary in other methodologies are eliminated. Using simple library amplification steps, a very small amount of starting material has been successfully used to profile genome-wide chromatin accessibility (9). Several studies have shown that 500-50,000 nuclei are sufficient for NGS analyses, considerably less than the nuclei required for other methods (9, 11). Thus, the small amount of cell numbers needed and its high resolution for mapping open chromatin regions make ATAC-seq an excellent tool for genome accessibility profiling.

ATAC-seq has been used to profile the chromatin structure of various animal samples (11-17), but such analyses on plants remain limited (18-20). Unlike for animal samples in which cells can be used directly as input in the transposition reaction immediately after lysing cells with a gentle detergent, plant tissues must be physically disrupted to release the cell contents from the rigid cell walls (18-21). Consequently, different tissues and plant species require individualized protocol optimization, making utilization of ATAC-seq in plant research more challenging.

Maize (*Zea mays*) is one of the most important crops globally, being widely consumed as staple food and livestock feed, as well as for its industrial uses. The maize genome is relatively large (∼2.3 Gbp) and has a complex organization of interspersed genes that are separated by transposable elements (22). In fact, 85% of genes are positioned within 1 kb of transposons (23). How maize cells coordinate this complicated chromatin structure with the transcription program in different tissues and developmental stages is a fascinating biological question. Many studies have explored massive transcriptional variation in different maize tissues and among different maize inbred lines (24-26). Interestingly, epigenetic analyses—such as DNA methylation, siRNA profiling, nucleosome occupancy and histone modification profiles—have suggested that epigenetics plays regulatory roles in gene expression and various aspects of maize development (27-33). Its genome complexity and intricate interplay between different levels of epigenetic control make maize an interesting model. Although the maize genome has been sequenced and related epigenetic features are being actively studied, its chromatin accessibility profile remains largely unknown. In this study, we describe the first ATAC-seq protocol for maize by testing several key steps. Unlike previously described methods (21), we found that liquid nitrogen-based homogenization is not suitable for maize ATAC-seq analysis, so we instead implement manual chopping for nucleus extraction. Importantly, we demonstrate that purification of maize nuclei to eliminate organellar DNA can be achieved without the need for cell sorting, and only a standard bench-top centrifuge is required for sample preparation. We also suggest the required parameters of library quality that enables the assessment of maize ATAC-seq libraries prior to sequencing. Finally, our sequence analyses validated that ATAC-seq can be used to analyze the maize genome.

## Results and Discussion

The most critical step in ATAC-seq protocols is the Tn5 transposition reaction during which the Tn5 transposase fragments and tags the DNA *in vivo* with unique Illumina library adaptors. Open chromatin regions have a higher probability of being targeted by Tn5, so several critical factors influence this step. First, high purity of nuclei is important. The Tn5 transposase not only targets genomic DNA, but also attacks the DNA of mitochondrial and chloroplast genomes, so it is better to have as few organelles as possible. In addition, an effective reaction requires Tn5 to be able to attack chromatin efficiently, so cell walls and cell debris should be eliminated. Second, during isolation, the native chromatin conformation must be preserved. Damaged nuclei and disordered chromatin structure could result in distortive results. Last, the ratio of transposase to the number of nuclei should be optimized based on genome size and chromatin structures. After an effective and efficient transposition reaction (Fig 1B), the resulting fragments tagged with adapters are subjected to two rounds of PCR to generate an ATAC-seq library for sequencing. In the first round of PCR, these fragments are amplified only by five PCR cycles during which distinct sequencing barcodes are added (Fig 1D). A fraction of this first PCR product is then subjected to quantitative PCR (qPCR) to estimate the relative amount of successfully tagged DNA fragments (Fig 1E) and then to determine the optimal amplification cycle number for the second round of PCR (Fig 1F). Under favorable conditions, an ATAC-seq library is ready when it contains sufficient DNA with proper sequence complexity after the second PCR. Since NGS is relatively expensive, applying an appropriate method for assessing library quality before sequencing can potentially reduce the sequencing expense (Fig 1G).

**Figure 1.**
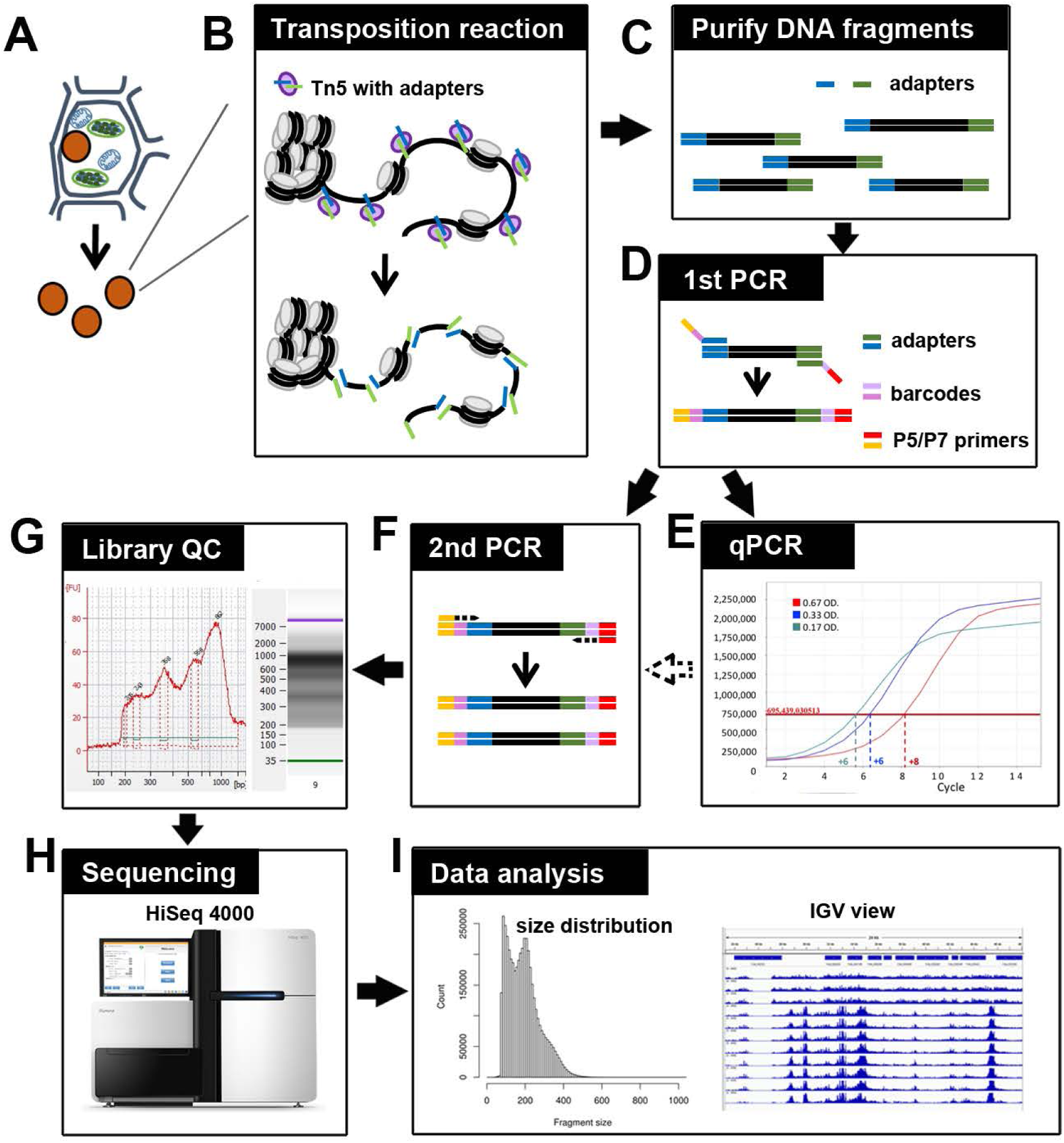
Graphic abstract: Flow chart of the procedure.

In this study, in order to provide a robust and efficient methodology for ATAC-seq analysis in maize, we focused on testing various parameters for key steps, including: (1) isolation of maize nuclei; (2) evaluation of optimal numbers of nuclei and effective Tn5 transposition; (3) library amplification conditions; and (4) library quality assessment. Finally, we validated our ATAC-seq results by comparing our data with nucleosome occupancy regions identified by MNase-seq (27). The ATAC-seq workflow is illustrated in Figure 1 and the detailed methodology is described in the supplemental information.

### Isolation of fresh nuclei from maize seedlings

A crucial requirement for generating a good ATAC-seq library is to use pure and intact nuclei (18). Moreover, since Tn5 not only targets nuclear DNA but also organellar DNA, contamination by organellar reads after sequencing can often account for up to 50% of the total sequencing reads (34). Removal of organelles prior to conducting the transposition reaction can thus greatly improve ATAC-seq efficiency. Two steps are usually implemented to isolate plant nuclei; breakage of plant cell walls to release nuclei, followed by organelle removal.

To break rigid plant cells, three different protocols are often applied, including grinding samples into a fine powder in liquid nitrogen, disrupting tissues using a polytron homogenizer, or slicing tissues using a sharp razor blade (35, 36). To establish which of these methods is best suited to obtaining good-quality intact nuclei for maize ATAC-seq, we tested all three methods. After filtering, nuclei were stained with DAPI for microscopic examination. As shown in Figure 2, the nuclei extracted by grinding in liquid nitrogen or disrupting by polytron homogenizer were of poor quality. Stringy mess or misconfigured nuclei suggested that cellular structures are destroyed and nuclei are also damaged. Chromatin released from broken nuclei exhibited irregular DAPI staining, often seen as scrambled mess with cell debris (Fig 2E), so the original chromatin structure likely has been altered. We found that it was difficult to conduct either of these two methods without damaging maize nuclei. However, when we adapted the protocol commonly used for protoplast preparation (37), in which maize fresh tissues are sliced and chopped with a sharp stainless steel razor blade, we observed high yields of round-shaped nuclei with distinct chromatin morphology (Figure 2C and D), in contrast to damaged nuclei isolated with two other methods (Figure 2A, B, and E). Interestingly, after collecting nuclei by centrifugation, we found that a large amount of starch grains appear in a pale white layer underneath a green layer. Starch grains do not stain with DAPI nor show autofluorescence (Figure 2F). To minimize starch grains in our samples, we found that a fine nylon paintbrush is very useful to separate the green layer hosting nuclei and most organelles from the underlying starch grain layer (see supplemental information for details).

**Figure 2.**
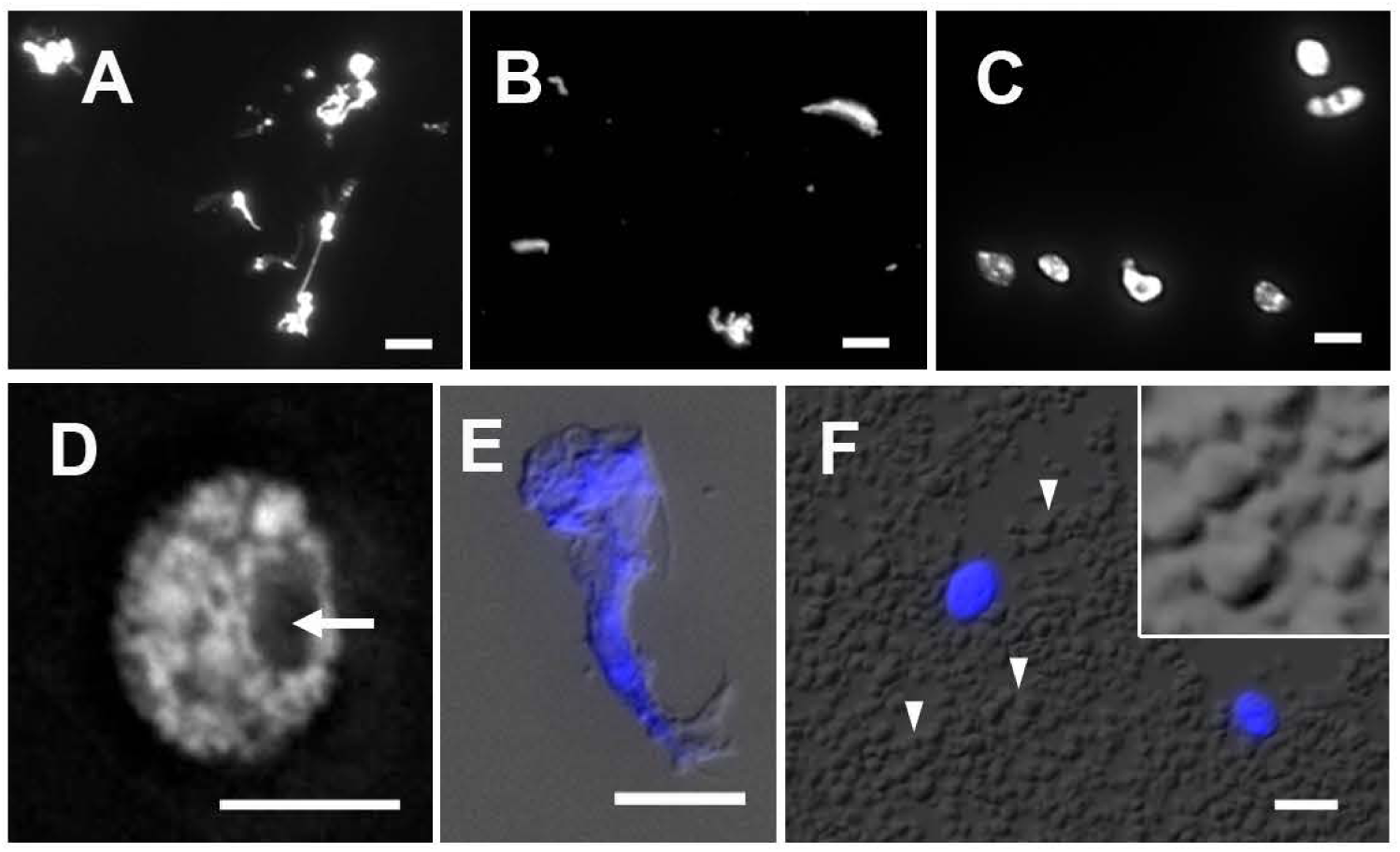
Nuclei stained with DAPI after extraction by (A) grinding in liquid nitrogen, (B) polytron homogenizer and (C) manual chopping. (D) Magnified image showing a round-shaped nucleus with distinct chromatin morphology and nucleolus (arrow) following the manual chopping method. (E) Magnified image of a cell debris showed distorted cell structure and scrambled chromatin with DAPI signals. (F) A large amount of starch grains (arrowheads) is observed in the crude homogenate. The magnified inset shows polyhedral starch grains. Scale bar represents 5 μm.

To eliminate organelles, an optimized concentration of non-ionic detergent, such as Triton X-100, is typically used to lyse organelle membranes, followed by purification steps. If flow cytometry is implemented, expressing a nuclear tagging fluorescent protein in plants will facilitate the efficiency of nuclei collection (18). Deal and Henikoff (38) developed a method, named the isolation of nuclei tagged in specific cell types (INTACT), which involves applying streptavidin-coated magnetic beads to isolate biotin-labeled nuclei extracted from plant samples. These methods enable purification of large numbers of nuclei, but require generation of transgenic plants. Traditionally, isolation of subcellular structures takes advantage of differential buoyant density, size and shape of subjects, which allows separation of subcellular compartments during centrifugation in high-viscosity media such as sucrose, mannitol, glycerol or Percoll (a colloidal silica). Production of transgenic maize lines is still a time-consuming process and ATAC-seq does not require a large number of nuclei, so we decided to adopt the Triton X-100 treatment, combined with a gradient centrifugation for nuclei purification. We first tested different concentrations (0.5-2%) of Triton X-100 to burst chloroplasts and adapted a protocol that uses a combination of sucrose and Percoll for density-gradient separations of nuclei (35). Figure 3 shows that both chloroplasts and nuclei remained intact after 0.5% Triton X-100 treatment, but most chloroplasts were lysed in 1% Triton X-100 buffer. However, numbers of intact nuclei declined when we used even higher concentrations of Triton X-100, suggesting that an optimal concentration for lysis of the most abundant organelles, i.e. chloroplasts, is 1% Triton X-100.

**Figure 3.**
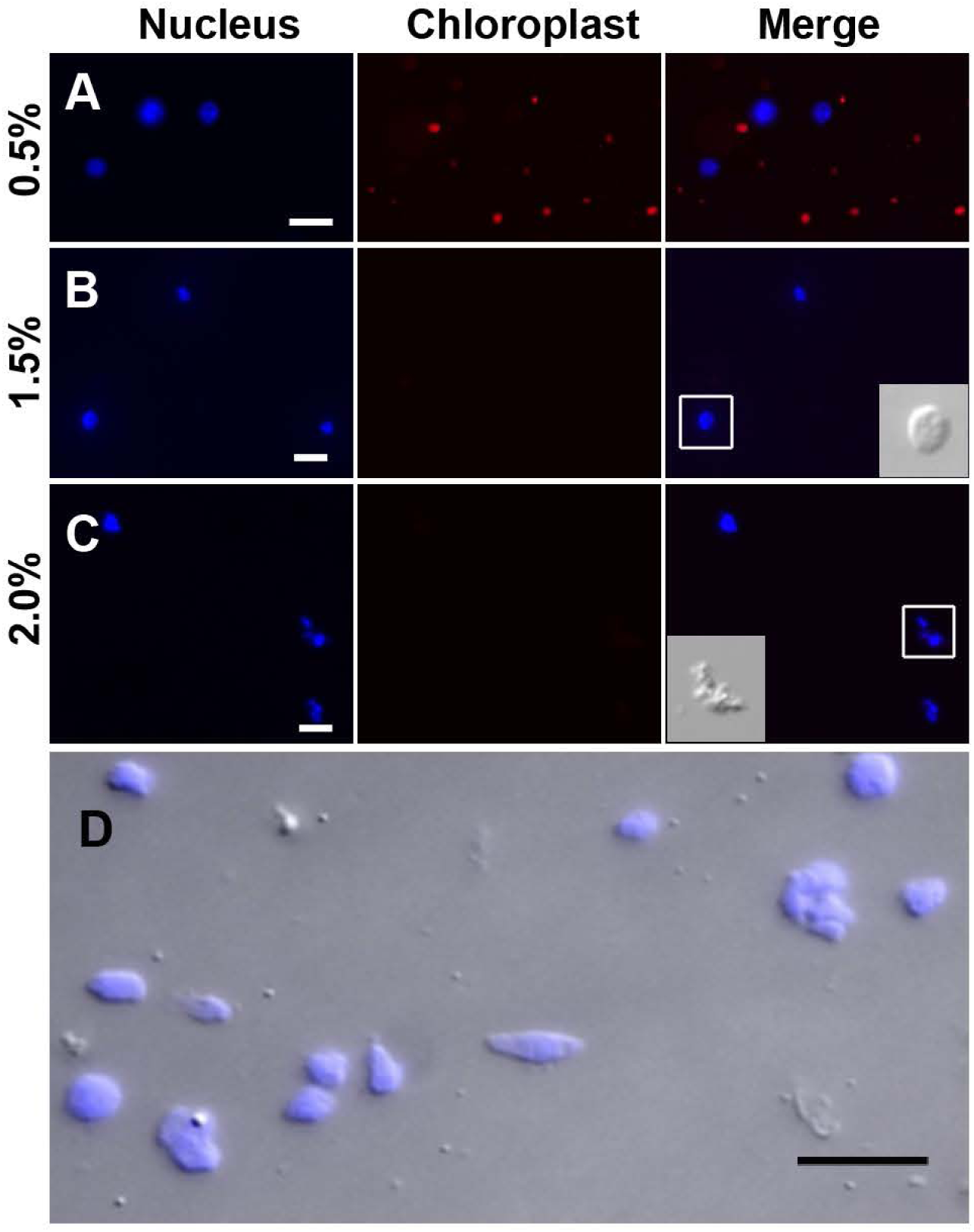
Maize nuclei isolation. Nucleus suspension after the detergent lysis step with (A) 0.5%, (B) 1.0%, (C) 2.0% Triton X-100. Nuclei are visualized with DAPI stain (blue) and chloroplasts exhibit strong red autofluorescence. Note that most chloroplasts are lysed in Triton X-100 above 1.0%. With higher Triton X-100 at 2.0%, nuclei integrity started to deteriorate. Insets in B and C are nuclei shown by DIC. Note that nucleus is disfigured in 2% Triton X-100. (D) Nuclei isolated by sucrose-Percoll gradient centrifugation remain intact (round-shaped). Scale bar represents 10 μm. Nuclei were stained with DAPI and detected using excitation wavelength 353 nm and emission wavelength 465 nm. Chloroplast autofluorescence was detected using 638 nm/646 nm (excitation/emission).

For sucrose-Percoll gradient separation, we tested 60% Percoll with 2.5 M sucrose as a cushion in a bench-top centrifuge (35) (Figure 4). After centrifuging at 1,000 x *g* for 15 min, we observed that the upper layer of supernatant was colored green due to chlorophyll content. A brownish-white layer was deposited on the sucrose-Percoll interface, which represents the nucleus fraction. Microscopic examination of this layer revealed intact round-shaped nuclei with a clear nucleolus (Figure 3D). We were able to collect approximately one hundred thousand pure nuclei from five seedlings using this approach of manual chopping, followed by 1% Triton X-100 treatment and 60% Percoll:2.5 M sucrose gradient separation.

**Figure 4.**
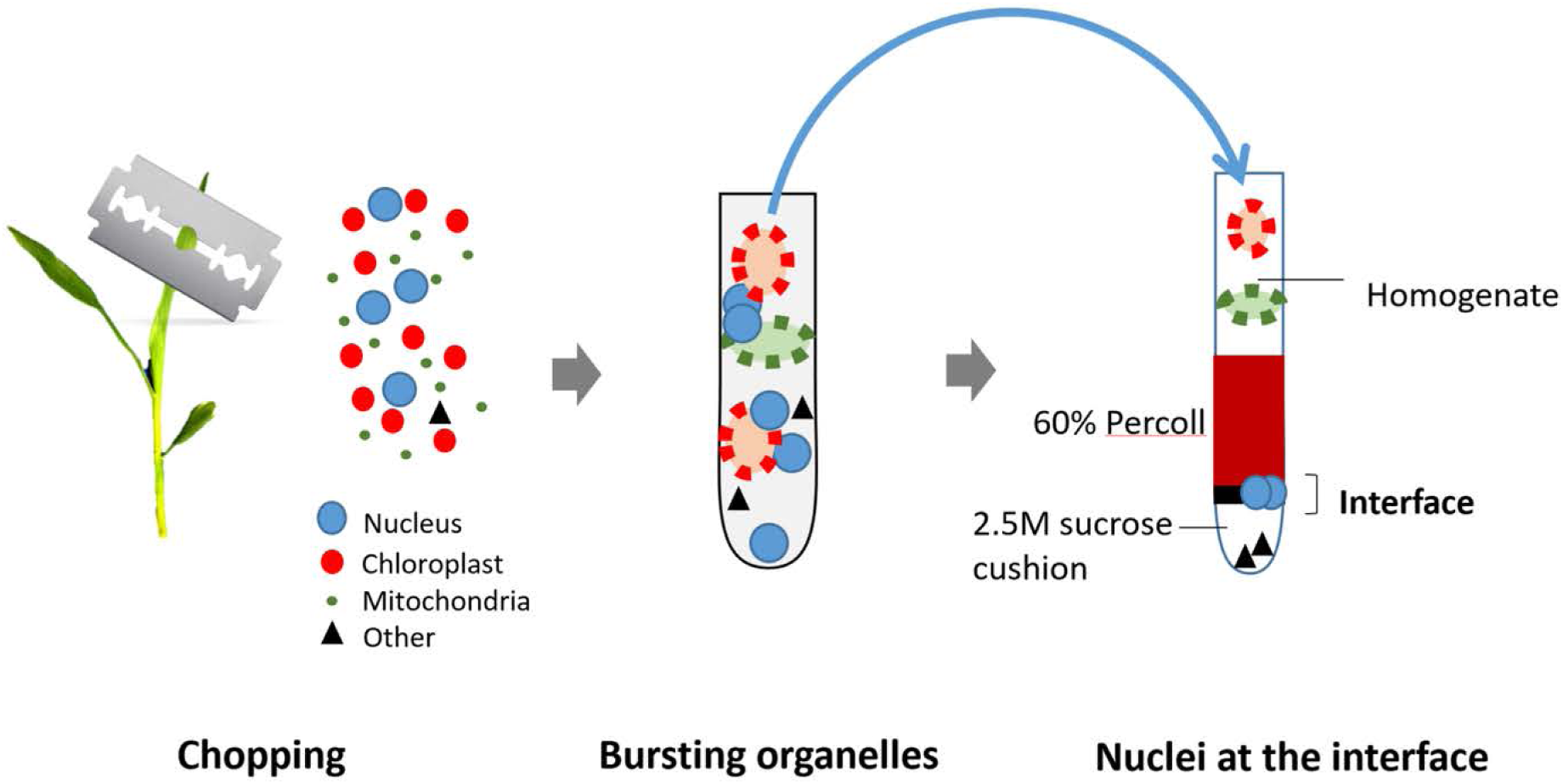
Nuclei isolation method. Fresh seedlings are chopped in a buffer with a sharp blade to release nuclei and organelles. Organelles are lysed using 1% Triton X-100, but nuclei remain intact. The crude exacts are then subjected to sucrose-Percoll gradient centrifugation. Nuclei are enriched at the interface between Percoll and the sucrose cushion. Isolated nuclei are transferred to a new tube and collected by centrifugation for the next transposition reaction.

### Optimization of nuclei number and Tn5 transposition efficiency

We used the Nextera DNA Library Preparation Kit (Illumina) to perform the Tn5 transposition reaction. According to the manufacturer’s manual, the input volume of nuclei suspension is limited to <5 μl, so we pelleted our isolated nuclei and resuspended them in a small amount of nucleus extraction buffer for the transposition reaction. To evaluate the appropriate number of maize nuclei for one reaction, we tested 500, 5,000, and 50,000 nuclei and used genomic DNA as a control. After the transposition reaction, DNA fragments were purified using PCR purification kit for the next amplification step. During the first round of PCR amplification, indexing barcodes are added onto each transposed DNA fragment. After five cycles of PCR, we paused the reaction and took a small aliquot (1 μl) of the PCR product to estimate the relative amount of effective DNA template using qPCR. Higher amounts of DNA template should be detected by qPCR if more transposition events have taken place, which ought to correlate with starting nuclei number in the reaction until saturation. As shown in Figure 5A, qPCR amplification plots from transposition reactions with 500 and 5,000 nuclei showed positive correlations with effective DNA template. However, reactions involving 50,000 nuclei did not produce more effective transposition events. In contrast, the naked genomic DNA exhibited a rapid amplification curve, suggesting that free DNA is much more easily targeted by the Tn5 reaction. Moreover, we tested whether using frozen nuclei could enhance transposition events (39). We froze approximately 2,500 purified nuclei at −20 °C for 30 min before proceeding with the transposition reaction. In comparison with 2,500 fresh (non-frozen) nuclei, qPCR analysis showed more transposed DNA fragments in the frozen sample, even exceeding the numbers generated in the reaction using 50,000 nuclei (Figure 5A). In addition, we tested whether a mild detergent in the transposition reaction could increase nuclear permeability for Tn5 transposition. Our results showed that SDS, CHAPS and NP40 did not result in significantly improved outcomes (Figure 5B).

**Figure 5.**
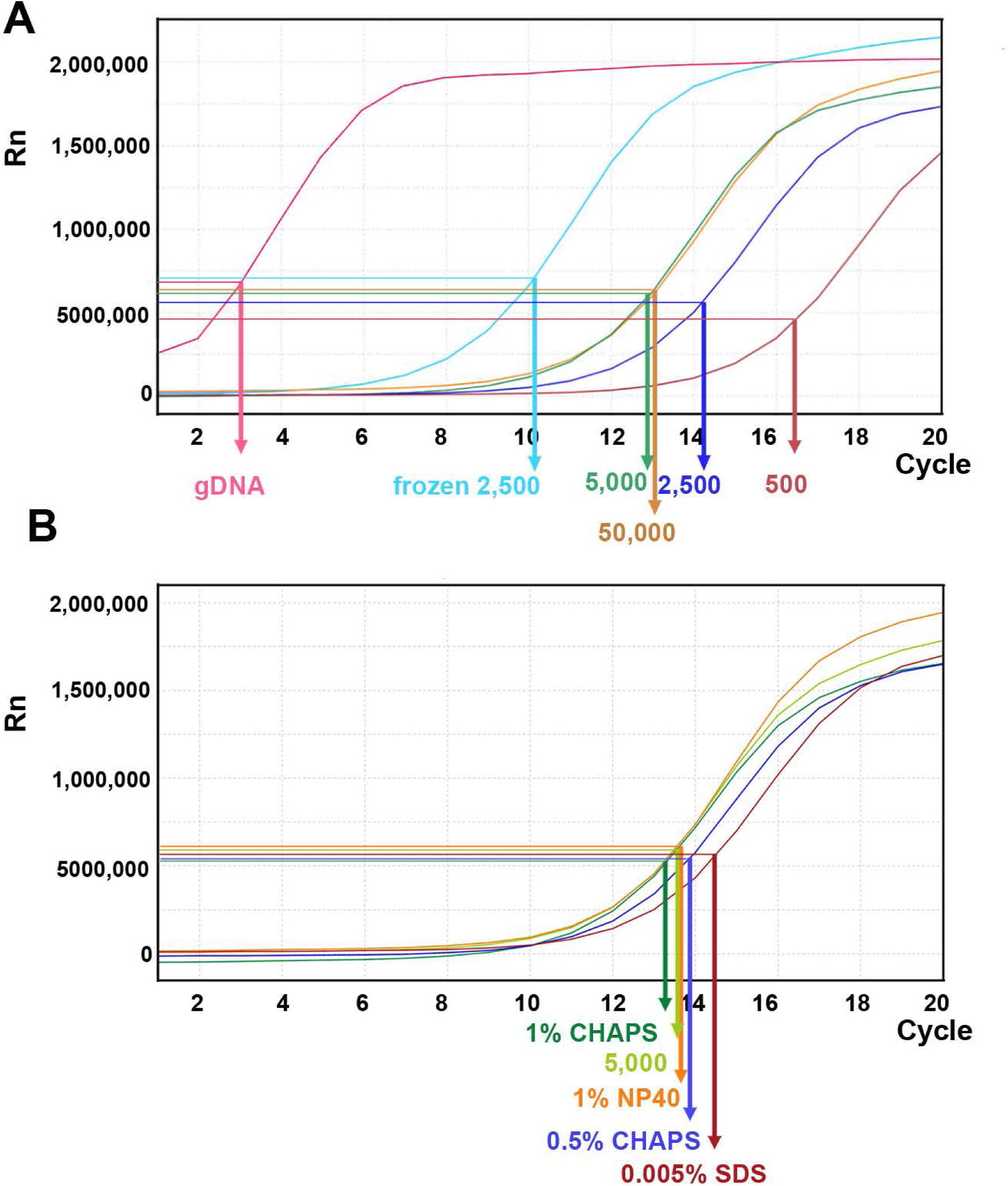
Linear amplification plot of quantitative PCR. (A) Quantitative PCR analysis for different transposition reactions. The linear amplification plot versus PCR cycles showed that reactions with fresh 500, 2,500, and 5,000 nuclei exhibited a positive correlation between cell numbers and amount of DNA template. However, the reaction with 50,000 nuclei did not exhibit more improvement. If 2,500 frozen nuclei were used for an input, the amplification plot showed a significant increase of DNA template amount. The genomic DNA (gDNA) severs as a control for transposition reaction. The arrows indicate the N cycle number of each sample corresponding to one-third of the maximum signal intensity. (B) Quantitative PCR analysis for transposition reactions with different mild detergent using 5,000 fresh nuclei.

### Determination of PCR cycle number for amplifying the NGS library

The first round of PCR with primers containing specific barcode and adapter sequences amplified those valid Tn5 transposition events after only five cycles (Figure 1). By qPCR analysis of a small aliquot from the first round of PCR, we estimated the relative amount of transposed DNA fragments, which can then be used to calculate the additional number of cycles (i.e. “N cycles”) necessary to amplify sufficient DNA for NGS. In the original protocol, 5 μl from the first PCR reaction was subject to qPCR analysis (9, 40). The “N” corresponds to the cycle number of qPCR when fluorescent intensity of PCR products reaches one-quarter to one-third of the maximum value, established by plotting linear Rn versus cycle number (Figure 5). It has been suggested that library complexity may become a concern if the total number of amplified cycles (5+N) is greater than 11 (9, 10, 41). In our modified protocol, only 1 μl was subject to qPCR analysis, so we consider that one fifth of DNA template was amplified in our qPCR analysis, which could roughly require two more cycles for teaching the same level of amplification in the original protocol. Thus, we expect to have a higher “N” in our experiments.

For our libraries of 2,500 and 5,000 fresh maize nuclei, we estimated N cycles of 14 and 13, respectively, based on the cycle number reaching one-third of the maximum value in our qPCR plots (Table 1, Figure 5). The fact that these N cycle numbers are even higher than we expected may suggest that the number of tagged fragments in maize is lower than achieved for other organisms, which may reflect that the maize genome contains a smaller proportion of open chromatin (18, 20, 21). Indeed, previous study showed that only a small portion (<1%) of the maize genome resides in open chromatin (27). To avoid excessive duplication and PCR bias, we tested the secondary PCR using a reduced number of additional cycles (four fewer than initially estimated). Accordingly, we generated sequencing libraries using 10 and 9 cycles for the 2,500 and 5,000 nuclei samples, respectively, so that the DNA concentrations of the libraries were just sufficient for sequencing but limited excessive duplicates (Table 1).

**Table 1.**
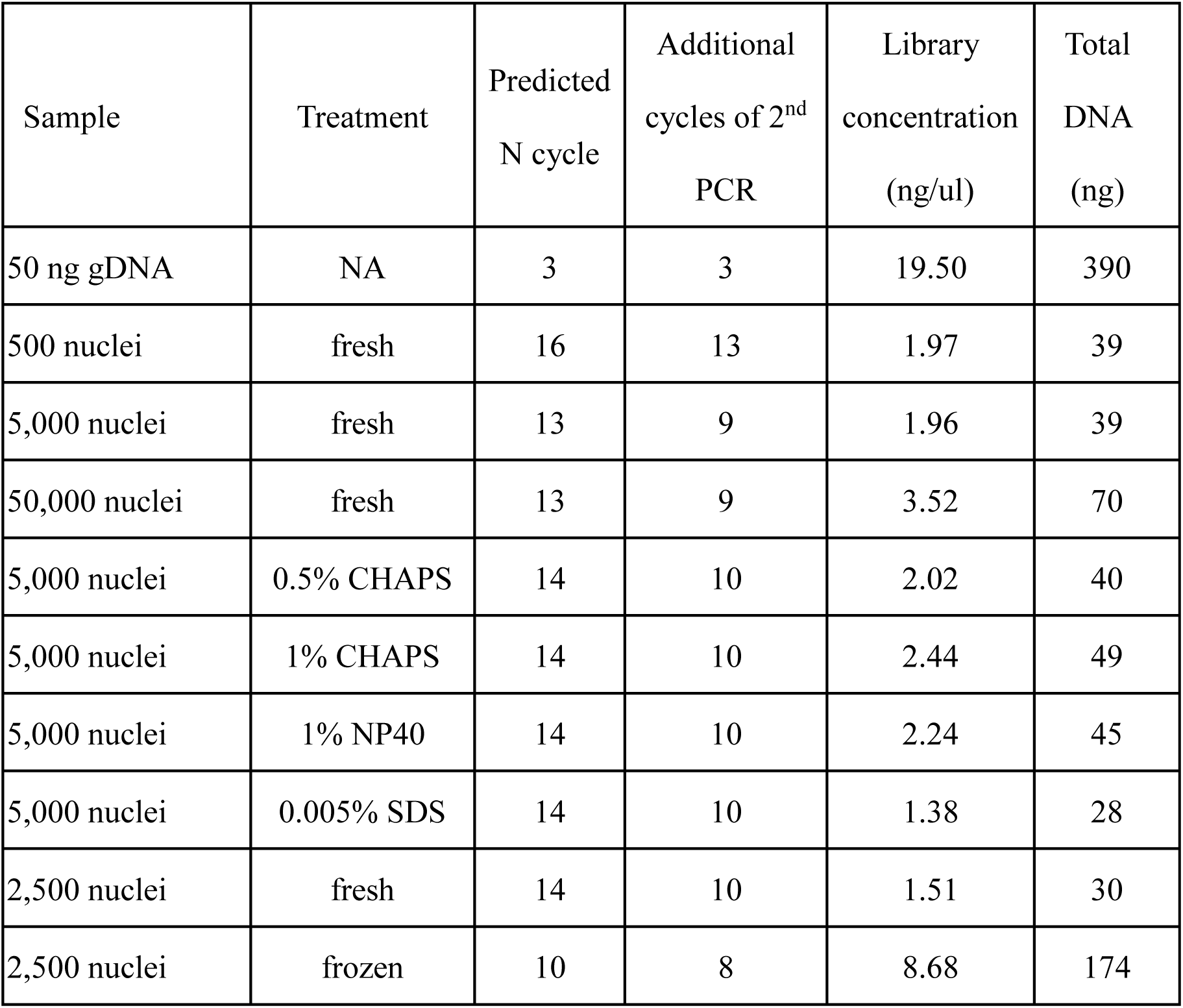
PCR cycles and DNA concentrations of ATAC-seq libraries.

### Library quality assessment

After the second round of PCR using sequencing primers, we quantitatively assessed resulting library output using a Qubit High Sensitivity Assay Kit (Thermo Fisher Scientific) and conducted quality evaluation by Bioanalyzer. This process allows us selecting libraries of good quality for further NGS investigation. As illustrated in Figure 1, Tn5 may insert adjacent to nucleosomes in open chromatin regions, which could give rise to transposed DNA fragments with lengths corresponding to mono- or multiple nucleosomes. Thus, a successful ATAC-seq library should exhibit a pattern of fragment size periodicity with intervals of around 200 bp. Additionally, transposed DNA fragments can result from two insertions in a non-nucleosomal DNA region. In contrast, if chromatin structures are damaged or perturbed, random transposition can result in a library of DNA fragments of various lengths lacking a consistent periodicity. Our Bioanalyzer analysis showed that using 2,500 or 5,000 fresh maize nuclei as input resulted in the distribution of fragment sizes adopting the expected ATAC-seq characteristics, i.e. size intervals of ∼200 bp (Figure 6B and 6C). Moreover, compared to the 2,500 nuclei sample, fragment sizes of the 5,000 nuclei sample were slightly shifted towards larger sizes, indicating that 5,000 nuclei may be approaching the upper limit of input quantity. In contrast, the reaction using 500 nuclei exhibited less a pronounced periodicity of peak (Figure 6A).

**Figure 6.**
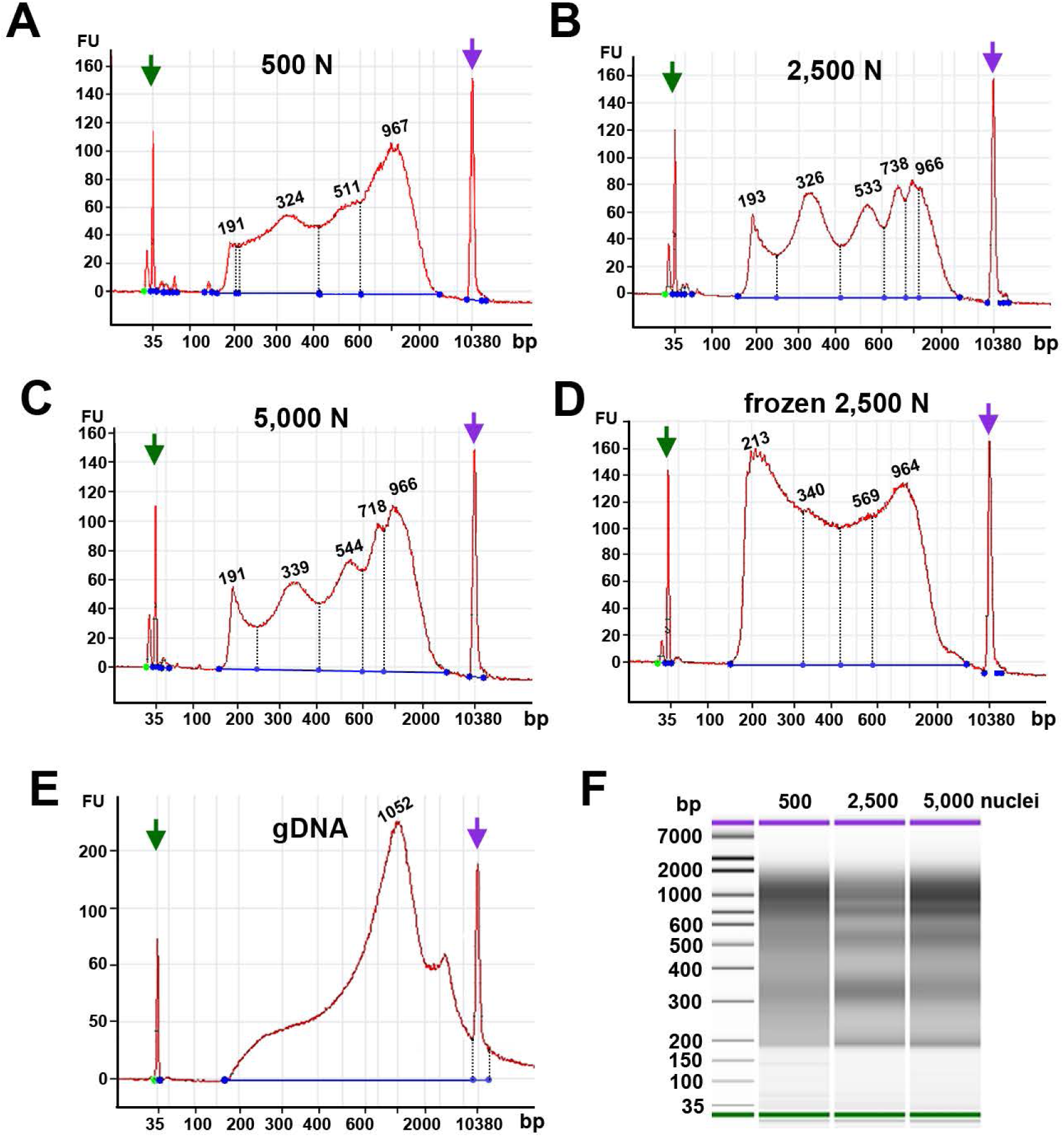
Fragment size distribution of ATAC-Seq libraries determined by an Agilent 2100 Bioanalyzer. The X-axis represents the product size in bp and the Y-axis is the arbitrary fluorescence intensity (FU). (A-E) Fragment size distribution plots of samples with (A) fresh 500 nuclei (500 N), (B) fresh 2,500 nuclei (2,500 N), (C) fresh 50,00 nuclei (5,000 N), (D) frozen 2,500 nuclei, and (E) 50 ng genomic DNA (gDNA). Numbers indicate fragment sizes of peaks, which may represent mono- or di- nucleosomes or more in 2,500 and 5,000-nuclei samples. Purple and green arrows represent upper and lower size markers, respectively. (F) A gel-like picture of fragment size distribution of ATAC-seq libraries made from 500, 2500, and 5,000 fresh nuclei.

Although our qPCR results suggested that frozen nuclei result in more efficient Tn5 transposition (Figure 5A), our Bioanalyzer data revealed that the frozen nuclei sample lacked a clear pattern of periodicity in fragment sizes (Figure 6D), suggesting that the chromatin structure may be disrupted by freezing or thawing. This outcome again supports that grinding maize samples in liquid nitrogen may not be suitable for maize ATAC-seq experiments. Similarly, Bioanalyzer data indicated that naked genomic DNA resulted in fragments with no size periodicity (Figure 6E). Taken together, these results suggest that 2,500 and 5,000 fresh maize nuclei represent a good starting material for ATAC-seq. In addition, maize ATAC-seq libraries can be generated successfully with an N cycle number lower than typically recommended, as long as the fragment length distribution indicated by Bioanalyzer analysis exhibits the desired characteristic periodicity (see below).

### Validation of ATAC-seq

In order to assess the results of our ATAC-seq protocols, we sequenced three ATAC-seq libraries constructed from fresh 2,500 nuclei, 5,000 nuclei and genomic DNA isolated from maize seedlings as a control. We assessed our libraries using two methods: 1) by plotting histograms of ATAC-seq fragment size distribution to visualize the characteristic size periodicity of ATAC-seq libraries (10); and 2) by comparing our ATAC-seq data against maize MNase-seq data, which identifies nucleosome-rich reads representing “closed” chromatin regions.

As outlined above, high quality ATAC-seq data should present a clear fragment size periodicity of approximately 200 bp (9), mimicking nucleosome turns. To compare fragment size distributions among the three libraries, we aligned the paired-end reads to the maize reference genome (AGPv4) using Bowtie2 (42), and then plotted the frequency of reads against the fragment size. As shown in Figures 7A and 7B, the libraries generated from 2,500 and 5,000 nuclei exhibited clear patterns of periodicity, indicating that both represent successful ATAC-seq libraries. In contrast, the library generated from naked genomic DNA (gDNA) showed no distinct size periodicity (Figure 7C), indicating that the Tn5 transposase had randomly inserted into genomic regions.

**Figure 7.**
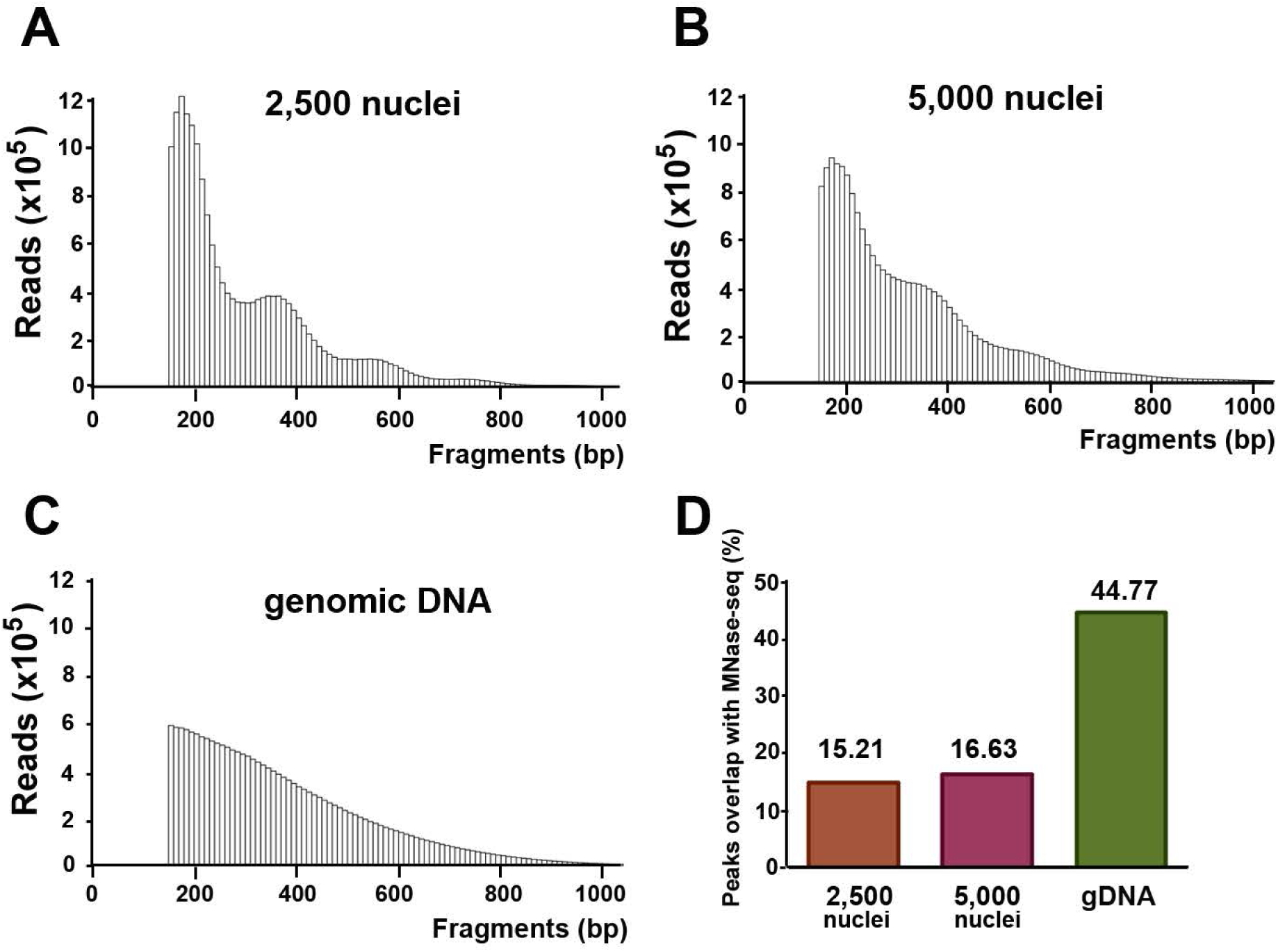
Bioinformatics analyses of ATAC-seq library reads. (A) Fragment size distribution of the 2,500-nuclei library. (B) Fragment size distribution of the 5,000-nuclei library. (C) Fragment size distribution of the gDNA control. (D) The percentage of overlapping peaks between ATAC-seq and MNase-seq datasets.

MNase-seq utilizes an endo-exonuclease that digests linker DNA between nucleosomes and unprotected DNA, revealing the inaccessible nucleosome-rich chromatin regions. Therefore, high quality ATAC-seq results should exhibit little overlap with the regions identified by MNase-seq. To examine the level of overlap between our ATAC-seq data and the available maize MNase-seq data derived from maize root samples (accession number SRR2542694; NCBI Sequence Read Archive), we conducted peak calling using MACS2 (43). Peaks from our ATAC-seq and MNase-seq overlapped greater than 80% were defined as overlapping peaks. With this criteria, only ∼ 15% of ATAC-seq peaks from both the 2,500 nuclei and 5,000 nuclei samples overlapped with those of MNase-seq. Thus, 85% of closed regions previously identified by MNase-seq analysis did not overlap with the open regions identified by our ATAC-seq. In contrast, peaks from the control gDNA sample presented 45% overlap between both approaches. Since the MNase-seq data was derived from 9-day-old root tissue, we do not expect to observe a complete exclusive pattern between our ATAC-seq and MNase-seq peaks. Overall, the ATAC-seq libraries we generated from both the 2,500 nuclei and 5,000 nuclei samples exhibited an appropriate size periodicity and limited overlap with closed chromatin regions identified by MNase-seq, indicating that the protocol described herein for generating ATAC-seq libraries is effective for identifying open chromatin regions in maize.

Our protocol employs a gentle nucleus extraction method to maintain nuclear integrity and, consequently, it faithfully retains chromatin structure. We conducted qPCR and Bioanalyzer analyses on our libraries, and only selected libraries for subsequent NGS that underwent minimal qPCR cycles and those that exhibited characteristic fragment size periodicity. These pre-screening approaches can eliminate unnecessary sequencing costs arising from poor quality libraries (Figures 6 and 7). Lastly, sequencing analyses using a Bioanalyzer validated the effectiveness of our protocol for preparing maize ATAC-seq libraries to profile the chromatin landscape of this important plant species.

### Conclusion

Chromatin accessibility is associated with gene expression, so it can influence phenotypes such as agricultural traits. Thus, a better understanding of chromatin architecture will provide great opportunities for crop improvement. The newly developed method for chromatin profiling, ATAC-seq, has been employed to investigate chromatin accessibility in human and many other animal species. However, there have been few reports of successful application of this chromatin profiling method in plants. In this study, we modified an original ATAC-seq protocol and applied it to the maize genome. First, we gently chopped fresh maize leaf tissue and then separated the nuclei using Percoll-sucrose gradient centrifugation. The isolated nuclei were treated with a transposase to fragment and tag accessible genome regions. The resulting fragments were subjected to two rounds of PCR to generate an ATAC-seq library. The qPCR analysis allows estimating the relative amount of valid transposition events, and determining the cycle number of the second PCR to amplify just sufficient DNA library. We assessed library quality using a Bioanalyzer prior to NGS, with distinct size peak periodicity of 200 bp indicating good quality. Computational analysis of fragment size distributions revealed a pattern of periodicity that is characteristic of ATAC-seq libraries. We found that 85% percent of closed regions previously identified by MNase-seq analysis did not overlap with the open regions established using our ATAC-seq approach, suggesting that the ATAC-seq library preparation procedure described herein is effective for identifying open chromatin regions of the maize genome.

## Material and methods

### Maize material and growth conditions

Maize seeds of the inbred line B73 were germinated with potting mix and grown in a growth chamber at 25 °C. Fresh aboveground tissue of 10-day-old plants after germination was collected immediately before the nuclear extraction procedure.

### Crude nuclear extraction

One seedling was homogenized by mortar and pestle in liquid nitrogen or by Polytron homogenizer, or by fine-chopping using a sharp razor blade with 1 ml cold NEB buffer. For the mortar and pestle method, plant tissue was first flash-frozen in liquid nitrogen and ground into fine powder. The pulverized tissue was then mixed in cold NEB buffer to release cell components. For the polytron method, fresh tissue was cut into smaller pieces and lacerated by a polytron (Omni TH motor with soft tissue OmniTips, Omni International Inc.) in cold NEB buffer. The setting of the polytron was 30 sec at mode ‘3’ and 1 min at mode ‘2’, followed by another 1 min at mode ‘1’. For chopping method, briefly, tissues were placed on parafilm and finely sliced with a sharp razor blade in cold NEB buffer. The slurry was filtered through 4 layers of Miracloth and a 20-μm CellTrics. Nuclei extracted by three different methods were gently pelleted by centrifugation at 800 xg for 10 min at 4°C.

In addition, while developing the chopping method, different concentrations of Triton X-100 (0.5%, 1.0% and 2.0%) were tested to determine the optimal amount of detergent to burst chloroplasts whereas nuclei remain intact. To assess the nuclear integrity and removal efficiency of chloroplasts, the suspension solutions were stained with DAPI (final concentration: 0.3 μg/ml) and observed with microscopy using excitation wavelength at 353 nm and emission wavelength at 465 nm for DAPI, and 638 nm/646 nm (excitation/emission) for chlorophyll (autofluorescence).

### Sucrose-Percoll gradient centrifugation

The Sucrose-Percoll gradient was used to further enrich nuclei in the Percoll-sucrose interface as described (35). Briefly, 400 μl 60% Percoll in ORB (as the ‘Percoll’ layer) was laid at the bottom of a 1.5-ml microcentrifuge tube. With a long gel-loading pipette tip, 200 ul SCB buffer (as the ‘Sucrose’ layer) was then carefully placed underneath the Percoll layer to ensure a clear demarcation between the two layers. The nucleus suspension was loaded on top of the Percoll-sucrose gradient buffer, and centrifuged at 1,000 xg for 15 min at 4°C.

### Tn5 transposition and library construction

The transposition and library construction steps are briefly described here. The detailed product information and experimental steps are indicated in the supplemental information. The Nextera DNA Library Preparation Kit (cat. FC-121-1031, Illumina) was used to perform the transposition according to the manufacturer’s manual. In addition to nuclear samples, 50 ng genomic DNA (gDNA), equivalent to 5,000-10,000 maize cells, was used as a negative control input. A brief description of the procedure is as follows. The nucleus pellet was suspended in the transposition reaction mix and incubated at 37°C for 30 min in an Eppendorf ThermoMixer Comfort (Eppendorf) with agitation at 1000 rpm for 2 min and resting for 2 min. The transposed DNA fragments were purified immediately after with a MinElute PCR Purification Kit (Cat. 28006, Qiagen).

After sequencing library was amplified as described in supplemental information, the library quality was preliminarily assessed prior to sequencing based on the total DNA concentration, the secondary qPCR cycle number and the Bioanalyzer pattern. Good quality libraries indicated by higher DNA concentrations, low secondary qPCR cycles and periodic BioAnalyzer patterns were selected to be sequenced by an Illumina platform HiSeq 4000 for 150-bp paired-end sequencing to generate 30 million raw reads for each maize library.

### Bioinformatic analysis of ATAC-Seq and MNase-seq

The ATAC-seq reads were aligned to the maize reference genome (AGPv4) using Bowtie2 (42), followed by the peak calling using MACS2 (43). The MNase-seq reads derived from maize root samples (accession number SRR2542694; NCBI Sequence Read Archive GSE49178), were processed in the same manner.

## Abbreviations

ATAC-seq: Assay for transposase accessible chromatin with high-throughput sequencing
NGS: Next generation sequencing
MNase: Micrococcal nuclease
DHSs: DNase I hypersensitive sites
qPCR: quantitative PCR
INTACT: isolation of nuclei tagged in specific cell types
gDNA: genomic DNA

## Accession Numbers

All sequence data can be downloaded from the NCBI Gene Expression Omnibus under accession number GSE120491.

## Availability of data and materials

All data generated or analyzed during the study are included in this published article.

## Acknowledgements

We thank the IPMB Genomic Technology Core Facility at Academia Sinica for their assistance with Bioanalyzer.

## Authors’ contributions

YJL and PC performed experiments and wrote the manuscript. JHL analyzed NGS results. PYC and CJRW designed experiments and wrote the manuscript.

