## Supplemental Information for "Assessing chromatin accessibility in maize using ATAC-seq"

**Basic Protocol 1: Isolation of fresh nuclei from maize seedlings**

***Materials***

10-day-old maize seedlings
Long gel-loading pipette tips (part number 1034-960-008, Labcon)
Percoll (P7828, Sigma-Aldrich)
Double edged stainless steel razor blade (Electron Microscopy Sciences, part number: 72000)
Glass Petri dish
Parafilm
MiraCloth
CellTrics cell strainers with 50 µm mesh (Sysmex Partec)
4',6-diamidino-2-phenylindole (DAPI) stain working solution (see recipe)
Nucleus Extraction Buffer (NEB) (see recipe)
Organelle Removal Buffer (ORB) (see recipe)
Sucrose Cushion Buffer (SCB) (see recipe)
Triton X-100 (20% (v/v)
Hemocytometer
1-ml and 2-ml Eppendorf low-binding tubes
Low-binding pipette tips
Fine nylon paintbrush (size 0)
cOmplete™, EDTA-free Protease Inhibitor Cocktail (part number: 04693159001; Roche)

***Nuclei isolation***

1. Cool the centrifuge to 4°C. Keep reagents and solutions on ice throughout the process.

2. Prepare the Percoll-Sucrose gradient buffer. Mix 400 µl ORB with 600 µl Percoll to make 60% Percoll. Add 400 µl 60% Percoll to the bottom of a new 1.5-ml microcentrifuge tube. Use a long gel-loading pipette tip to carefully lay 200 µl SCB underneath the Percoll layer, ensuring there is a clear demarcation between the two layers. Keep on ice. Perform this step at least 1 hour before using the Percoll-sucrose gradient

   *Note: We found that one-hour equilibration of the Percoll-Sucrose gradient buffer resulted in better nuclei recovery than immediate use..*

3. Place a piece of 5x5 $cm^2$ parafilm on a glass Petri dish on ice. Pipet 500 µl NEB onto the parafilm and chop above ground tissue from 10-day-old seedlings (approximately 5 seedlings) in NEB buffer with a double edged razor blade. Add more NEB buffer if needed until the chopped tissue has the appearance of a coarse slurry. Complete tissue chopping in 2-3 batches. Approximately 5 ml NEB is needed.

   *Note: Chopping with a sharp razor blade is a gentler method to release nuclei without damaging them compared to grinding in liquid nitrogen in our hands.*

4. Transfer the slurry to a small petri dish on ice. Incubate for 5 min with gentle agitation. Place 4 layers of Miracloth on a pre-chilled glass funnel. Filter the slurry through the

funnel into a 15-ml Falcon tube. Gently squeeze the remaining tissue against the funnel to extract more nuclei.

5. Filter this crude nuclei suspension through a 20-µm CellTrics twice to remove large debris. Divide the filtrate into 1.5-ml microcentrifuge tubes and centrifuge at 800 xg for 10 min at 4°C.

6. Gently remove the supernatant without disturbing the pellet. The pellet should comprise a pale-white layer below and a green layer on top.

   *Note: The white layer predominantly consists of starch grains, and the top layer contains nuclei, chloroplasts and other organelles.*

7. Add 200 µl ORB (with 1% Triton X-100) into each tube from step 6. Incubate for 2 mins so the pellet loosens. Use a fine nylon paintbrush to gently remove the green layer from the pellet, leaving the pale-white bottom layer undisturbed. Gently transfer the green pellets in ORB to a new tube by pipetting. Resuspend the green pellet material in the ORB using the paintbrush and by gentle pipetting.

   *Note: Chloroplasts are broken up by 1% Triton X-100 in the ORB buffer. If this leftover solution is centrifuged, the pellet will be white and lack the uppermost green layer as chlorophyll is released in the supernatant from broken chloroplasts. The pale-white lower layer is rich in starch grains.*

8. Filter this nuclei suspension through a 10-µm CellTrics twice to remove aggregate nuclei and debris.

9. Load approximately 400 µl of this green-colored nucleus suspension on the previously-prepared Percoll-sucrose gradient buffer. Centrifuge at 1,000 xg for 15 min at 4°C. Use another Percoll-Sucrose gradient, if more than 400 µl nuclei suspension solution is being used.

10. Remove the green-colored supernatant on the top. A brownish-white layer deposited on the Percoll/SCB interface should be observable, which represents the nuclei fraction. Carefully transfer this nuclei fraction (~ 200 ul) into a new 1.5-ml microcentrifuge tube.

    *Note: In practice, some solution from the Percoll-sucrose and SCB layers can be also taken up, together with nuclei fraction.*

11. Add 1× volume of NEB to dilute the nuclei fraction suspension. Centrifuge at 500 xg for approximately 10 min to pellet the nuclei.

    *Note: Check the pellet condition every 2 mins, and stop centrifugation when a small translucent pellet is first seen. We found that excessive centrifugation results in nuclear damage. The pellet is translucent, different from the pale-white color of the starch grain pellet.*

12. Dissolve the pellet in 20 µl NEB to produce the final isolated nuclei solution.

13. Take 2 µl of the isolated nuclei solution and dilute in NEB. Stain with DAPI (final concentration: 0.3 µg/ml). Examine the nuclei integrity and calculate the nucleus density in the isolated nuclei suspension from Step 12 with a hemocytometer.

    *Note: The expected nuclei concentration from the Step 12 is approximately 5-20 million/ml.*

14. Transfer a volume equivalent to 2,500 – 5,000 nuclei to a new 1.5-ml low-binding tube.

    *Note: The input volume of nuclei for the transposition reaction must be < 5 ul.*

**Basic Protocol 2: Tn5 Transposition, Library Preparation and Sequencing**

***Materials***

Purified intact nuclei
Qubit dsDNA BR Assay Kit (Part Number: Q32853, Thermo Fisher Scientific)
Nextera® DNA Library Preparation Kit (Part Number: FC-121-1031, Illumina)
1.7ml Microtube, Clear, Maxymum Recovery (Part Number: MCT-175-L-C, Axygen)
NEBNext High-Fidelity 2x PCR Master Mix (Part Number: M0541S, New England Labs)
Nextera Index Kit (Part Number: FC-121-1011, illumina)
MinElute PCR Purification Kit (Part Number: 28006, Qiagen)
Agencourt AMPure XP (Part Number: A63881, Beckman Coulter)
80% Ethanol (Part Number: 1.00983.2500, Merck)
UltraPure DNase/RNase-Free Distilled Water (Part Number: 10977-015, Thermo Fisher Scientific)
Buffer EB (Part Number: 19086, Qiagen)
Library Quantification Kit Illumina® platforms ABI PrismTM qPCR Master Mix (Part Number: KK4835, KAPA Biosystems)
Qubit dsDNA High Sensitivity Assay Kit (Part Number: Q32854, Thermo Fisher Scientific)
High Sensitivity DNA Kit (Part Number: 5067-4626, Agilent Technologies)
Transposition reaction mix (see recipe)
First PCR amplification reagents (see recipe)
qPCR reagents (see recipe)
Secondary PCR amplification reagents (see recipe)

***Equipment***

Thermomixer Comfort (Catalog number: 5355, Eppendorf)
T100™ Thermal Cycler (Catalog number: 186-1096, BIO-RAD)
Centrifuge: MiniSpin Plus (Catalog number: 5453, Eppendorf)
Applied Biosystems QuantStudio™ 12K Flex Real-Time PCR System (Catalog number: 4470050, Thermo Fisher Scientific)
DYNAL MPC-S magnetic stand (Catalog number: A13346, Thermo Fisher Scientific)
Qubit™ 2.0 (Catalog number: Q32866, Thermo Fisher Scientific)
Agilent 2100 Bioanalyzer Instrument (Catalog number: G2939BA, Agilent Technologies)

***Tn5 Transposition***

The Nextera DNA Library Preparation Kit (FC-121-1031, Illumina) was used to perform the transposition reaction according to the manufacturer's manual. In addition to nuclear samples, 50 ng genomic DNA (gDNA) was used as a negative control input. A brief description of the procedure is as follows.

15. Prepare the transposition reaction mix (see recipe) and resuspend the nucleus pellet (~2,500 nuclei) in the transposition reaction mix.

    *Note: According to our experience, the optimal number of nuclei for one reaction is 2,500 – 5,000.*

16. Incubate the transposition reaction at 37°C for 30 min in an Eppendorf ThermoMixer Comfort with agitation at 1000 rpm for 2 min and resting for 2 min (i.e. occasionally mixing).

    *Note: Occasional gentle mixing may increase fragment yield.*

17. Immediately following transposition, purify the Tn5 transposed DNA fragments with a Qiagen MinElute PCR Purification Kit.

***Purify DNA fragments***

Use a Qiagen MinElute PCR Purification kit to purify the transposed sample according to the manufacturer's manual. A brief description of the procedure is as follows.

18. Add 5X volume of PB Buffer into the transposition reaction and vortex to mix.

    *Note: Check for the yellow color of PB Buffer. If the buffer turns orange or violet, the pH is too high. A small volume of 3M sodium acetate can be added to adjust pH before proceeding.*

19. Transfer the mix into a MinElute spin column provided by the Qiagen MinElute PCR Purification kit.

20. Centrifuge for 1 min at 13,000 rpm. Discard all the flow-through.

21. Add 750 µl of PE Buffer, and centrifuge for 1 min at 13,000 rpm. Discard the flow-through.

22. Centrifuge for 1 min at 13,000 rpm. Rotate the column by 180° and centrifuge for 1 min.

    *Note: This step removes any remaining ethanol in the column.*

23. Place the MinElute spin column in a new low binding tube.

24. Elute in a new tube with 12 µl of EB buffer (10 mM Tris, pH 8.5) and stand at room temperature for 1 min.

25. Centrifuging for 1 min at 13,000 rpm. Rotate the column 180° and centrifuge for 1 min.

26. Discard the column and store the purified DNA at -20°C if necessary.

    *Note: This is a convenient stopping point for the protocol.*

***Preparation of sequencing libraries***

This phase consists of five major steps. First, indexing barcodes are added onto each of the transposed DNA fragment by 5 cycles of PCR amplification ('the first PCR'). Second, a small aliquot of the product generated from the first PCR is subjected to quantitative PCR (qPCR) to assess amount of DNA template for determining the number of cycles in the secondary PCR. Third, these indexed fragments integrated in the First PCR are then further amplified with sequencing primers (i.e., 'the secondary PCR') to generate a sufficient amount of DNA for sequencing. Fourth, the resulting library from the secondary PCR is purified, and last, its quality is estimated by Bioanalyzer before being sequenced.

***First PCR amplification with indexing barcode primers***

Use Nextera Index Kit and NEBNext High-Fidelity 2x PCR Master Mix to amplify transposed DNA fragments. A brief description of the procedure is as follows.

27. To amplify transposed DNA fragments, combine the First PCR Amplification Reagent (see recipe) with 10 µl purified transposition product in a 0.2 ml PCR tube.

    *Note: Different index primers (barcodes) should be selected for different samples, so these samples can be pooled together for sequencing. After sequencing, these samples can be separated according to their barcodes.*

28. The thermal cycle for PCR amplification is as follows: 72°C for 5 min, 98°C for 30 sec, 5 cycles (98°C for 10 sec, 63°C for 30 sec, and 72 °C for 1 min).

29. Transfer the PCR product to a new microcentrifuge tube.

***Determining the cycle number for the secondary PCR by qPCR***

The KAPA Library Quantification Kit is used for the quantitative PCR. A brief description of the procedure is as follows.

30. Mix these qPCR reagents (see recipe) with 1 µl of the amplification product from the first PCR amplification.

31. Perform qPCR amplification using an Applied Biosystems QuantStudioTM 12K Flex Real-Time PCR System with the following conditions: 98°C for 5 min, 20 cycles (98°C for 10 sec, 63°C for 30 sec, and 72 °C for 1 min).

32. Calculate the required number of cycles (i.e., 'N' cycles) for the secondary PCR step by plotting linear Rn versus the cycle number. The N cycle number corresponds to the cycle number at which fluorescent intensity reaches one-third of the maximum value

***Secondary PCR amplification***

Use the NEBNext High-Fidelity 2x PCR Master Mix to perform the secondary amplification step.

33. Perform the secondary PCR with the remaining product from the first PCR, and its cycle number is determined by the previous qPCR (Step 32).

34. Mix the Secondary PCR Amplification reagents (see recipe) and the 40 μl of the first PCR product in a 0.2 ml PCR tube.

35. Perform the secondary PCR amplification as follows: 98°C for 30 sec, and N cycles of (98°C for 10 sec, 63°C for 30 sec, and 72 °C for 1 min). N cycles was determined by qPCR.

    *Note: Based on our experience, successful libraries can be generated using a cycle number lower than the determined by qPCR.*

***Purify the amplified library with AMPure XP beads***

Use AMPure XP kit to purify the secondary PCR amplified fragments. A brief description of the procedure is as follows.

36. Allow the AMPure XP beads to come to room temperature for at least 30 min. Vortex the AMPure XP Beads until they are well dispersed. Add 100 μl (1X) of well-mixed AMPure XP Beads into a new 1.7 ml tube.

37. Transfer each DNA library to a new 1.7 ml tube containing the AMPure XP Beads. Gently pipette the entire volume up and down 10 times or vortex gently to mix thoroughly.

38. Incubate the tubes at room temperature for 5 mins. Place the tubes on a magnetic stand at room temperature for 3 mins or until the liquid appears clear. Remove the supernatant from each tube.

    *Note: Some liquid may remain in each tube. Do not disturb the beads.*

39. Wash the pellet twice with 200 μl of 80% ethanol. Let the tubes stand at room temperature for 10 mins to dry.

40. Resuspend the dried pellet in each tube with 22.5 μl EB Buffer (see recipe). Vortex to mix thoroughly and incubate the tube at room temperature for 2 mins.

41. Place the tube on a magnetic stand at room temperature for 3 mins or until the liquid appears clear. Transfer all of the clear supernatant to a new 1.7 ml tube. The samples can be stored at -20°C.

***Checking library quality and sequencing***

42. Use Qubit High Sensitivity Assay Kit to determine the concentration of each ATAC-seq library.

43. Determine the fragment size distribution and peak pattern of each library using the Agilent Bioanalyzer 2100 system.

    *Note: A successful ATAC-seq library should exhibit a pattern of periodicity in fragment sizes with an interval of around 200 bp.*

44. We used an Illumina platform HiSeq 4000 for 150-bp paired-end sequencing to generate 30 million raw reads for each maize library.

    *Note: Other Illumina sequencing platforms can be used. Choose ones capable of paired-end sequencing with a minimal read length of 75 bp.*

### REAGENTS AND SOLUTIONS

#### For Basic Protocol 1:

Use sterilized Milli-Q water for all recipes and protocol steps.

***Nucleus Extraction Buffer (NEB) (10 ml)***

200 µl MOPS stock solution (1 M, pH 7.0) (see below)
80 µl NaCl stock solution (5 M)
300 µl KCl stock solution (3 M)
50 µl EDTA stock solution (500 mM)
50 µl EGTA stock solution (100 mM)
5 µl spermine stock solution (400 mM)
13 µl spermidine stock solution (400 mM)
200 µl protease inhibitor stock solution (see below)

Mix all stock solutions and bring the total volume to 10 ml with Milli-Q water. The buffer can be stored at 4 °C for 1 week.
Add protease inhibitor, spermine, and spermidine stock solutions right before use.

***1M MOPS stock solution (100 ml)***

Dissolve 20.93 g 3-(N-morpholino)propanesulfonic acid, 4-morpholinepropanesulfonic acid (MOPS) in 80 ml Milli-Q water.
Adjust the pH with 10N NaOH.
Autoclave and store at 4°C for a maximum of 3 months.

***Protease inhibitor stock solution***

Dissolve with vortexing one cOmplete™, EDTA-free protease inhibitor cocktail tablet in 1 ml Milli-Q water to make 50x stock solution with vortex. Store at -20C for up to 12 weeks.

***Organelle Removal Buffer (ORB) (10 ml)***

1 ml Sucrose stock solution (2.5 M)
100 µl Tris-HCl (1 M, pH8.0)
1 ml MgCl2 (100 mM)
500 µl Triton X-100 (20%)
200 µl protease inhibitor stock solution

Mix all stock solutions and Triton X-100.
Bring the total volume to 10 ml with Milli-Q water. The buffer can be stored at 4 °C for 1 week.
Add the protease inhibitor stock solution right before use.

***20% (v/v) Triton X-100 (100 ml)***

Add 20 ml Triton X-100 into 60 ml Milli-Q-purified water.
Stir for 30 min until fully mixed.

Store in the dark at room temperature for a maximum of 2 months.

***Sucrose Cushion Buffer (SCB) (10 ml)***

6.8 ml Sucrose stock solution (2.5 M)
100 µl Tris-HCl (1 M, pH8.0)
200 µl MgCl2 (100 mM)
50 µl Triton X-100 (20%)
200 µl protease inhibitor stock solution

Mix all stock solutions and Triton X-100.
Bring the total volume to 10 ml with Milli-Q water. The buffer can be stored at 4 °C for 1 week.
Add the protease inhibitor stock solution right before use.

***DAPI stock solution (5 ml)***

10 mg DAPI dilactate
Milli-Q-purified water

Dissolve 10 mg DAPI in 1 ml Milli-Q-purified water with vigorous shaking.
To make a working DAPI solution, dilute the stock solution 1000 x to 1 µg/ml. For long-term storage, the stock solution can be aliquoted and stored at -20 °C for at least six months.

**For Basic Protocol 2:**

Use nuclease-free water in all recipes and protocol steps.

***Transposition reaction mix (50 µl)***

20.5 µl Nuclease free water
25.0 µl 2 x TD reaction buffer
2.5 µl TDE1 (Nextera Tn5 Transposase)
2.0 µl Purified intact nuclei (2,500 nuclei) or 50 ng gDNA

***First PCR amplification reagents (50 µl)***

10.0 µl Nuclease free water
25.0 µl NEBNext High-Fidelity 2x PCR Master Mix
2.5 µl 25 µM Index 1 (i7)
2.5 µl 25 µM Index 2 (i5)
10.0 µl Transposed DNA

***qPCR reagents (20 µl)***

7 µl Nuclease free water
12 µl 2X KAPA SYBR® FAST qPCR Master Mix with 10X Primer Premix
1 µl first PCR product

***Secondary PCR Amplification reagents (100 µl)***

5 μl Nuclease free water
50 μl NEBNext High-Fidelity 2x PCR Master Mix
5 μl PCR Primer Cocktail
40 μl first PCR amplification product
